# Epigenetic compound screening uncovers small molecules for re-activation of latent HIV-1

**DOI:** 10.1101/2020.08.21.262311

**Authors:** Ariane Zutz, Lin Chen, Franziska Sippl, Christian Schölz

**Affiliations:** Max von Pettenkofer Institute & Gene Center, Virology, National Reference Center for Retroviruses, Faculty of Medicine, LMU München, Feodor-Lynen Str. 23, 81377 Munich, Germany

**Keywords:** epigenetic compound screen, human immunodeficiency virus (HIV), HIV-1, latency, latency reversing agents (LRA)

## Abstract

During infection with the human immunodeficiency virus type 1 (HIV-1), latent reservoirs are established, which circumvent full eradication of the virus by antiretroviral therapy (ART) and are the source for viral rebound after cessation of therapy. As these reservoirs are phenotypically undistinguishable from infected cells, current strategies aim to reactivate these reservoirs, followed by pharmaceutical and immunological destruction of the cells.

Here, we employed a simple and convenient cell-based reporter system, which enables sample handling under biosafety level (BSL)-1 conditions, to screen for compounds that were able to reactivate latent HIV-1. The assay showed a high dynamic signal range and reproducibility with an average Z-factor of 0.77, classifying the system as robust. The assay was used for high-throughput screening (HTS) of an epigenetic compound library in combination with titration and cell-toxicity studies and revealed several potential new latency reversing agents (LRAs). Further validation in well-known latency model systems verified earlier studies and identified two novel compounds with very high reactivation efficiency and low toxicity. Both drugs, namely N-hydroxy-4-(2-[(2-hydroxyethyl)(phenyl)amino]-2-oxoethyl)benzamide (HPOB) and 2',3'-difluoro-[1,1'-biphenyl]-4-carboxylic acid, 2-butylhydrazide (SR-4370), showed comparable performances to other already known LRAs, did not activate CD4^+^ T-cells or caused changes in the composition of PBMCs as shown by flow cytometry analyses.

Both compounds may represent an effective new treatment possibility for revocation of latency in HIV-1 infected individuals.

## Introduction

Approximately, 37.9 million people worldwide are living with HIV and about 65 % of them were accessing antiretroviral therapy (ART) (1). While ART effectively blocks viral replication, prevents further transmission, and results in partial reconstitution of the immune system and thus drastically reduces HIV-associated morbidity, it fails to eradicate the virus entirely. The reason for this is the establishment of transcriptionally silent and long-lived cell reservoirs during infection, which stay unaffected by ART and represent the source of viral rebound upon cessation of therapy (2–4). These reservoirs are generated early during acute infection (5) and despite a relatively low frequency of ∼1/10^6^ in resting CD4^+^ T-cells in treated individuals (6), they are very stable with a half-life of more than 3 years (7, 8). It is believed that either clonal expansion of latently infected cells itself (9) or a sustained low-level viral replication – especially in tissues with suboptimal concentrations of ART – is responsible for reservoir maintenance (10). Research exploring the nature of HIV-1 latency has accelerated in recent years, indicating that in particular epigenetic silencing of nucleosomal histones, the viral promoter region (long terminal repeat; LTR) as well as modulation of relevant transcription factors by posttranslational modifications (PTMs) play a key role in sustained viral persistence (reviewed in more detail in (11)). Thereby, especially de-acetylation events by histone deacetylases (HDACs) such as HDAC1 and HDAC2 (12), but also the transfer of methyl-groups by histone-lysine N-methyltransferases (13, 14) seem to be pivotal. Thus, novel strategies for the treatment of HIV have been employed, known as “shock and kill” approach, which focus on transcriptional reactivation of latent reservoirs by epigenetic-active drugs and subsequent prevented spreading and eradication by classical ART (15). In this context, HDAC inhibitors (HDACis) have been shown in a variety of experimental systems as well as in cells from HIV-infected individuals to be able to act as latency reversing agents (LRA) *in vitro* and *in vivo* (16). Furthermore, combinations of HDACis and protein kinase C agonists have been shortlisted for clinical trials (17–21). However, despite first encouraging results, clinical studies so far have yielded only limited success. For instance, vorinostat (also known as zolinza or suberoylanilide hydroxamic acid (SAHA)) a pan-HDACi as well as romidepsin (depsipeptide, istodax), a potent HDAC1 and HDAC2 inhibitor and valproic acid, targeting HDAC1-3, exhibited profound reactivation of HIV but also disadvantageous off-target effects. Moreover, none of the three compounds caused a significant reduction of the viral reservoir (22–26). The examples clearly demonstrate that despite several important qualities of these HDACis as LRA, there is still a high demand of further research to identify new – so far unappreciated – substances, which might be able to eradicate latent HIV reservoirs in infected individuals.

In this study we analyzed an unique epigenetic compound library, consisting of a collection of 380 substances with biological activity used for epigenetics research. Substances are structurally highly diverse, medicinally active, cell permeable, and include inhibitors of HDACs, sirtuins (SIRTs), lysine demethylases, histone acetyltransferases (HATs), DNA methyltransferase (Dnmts), and sirtuin activators (**Supplemental Figure 1**). Each compound was tested for its concentration-dependent effectivity in latency reversal as well as for its cellular toxicity, using an affordable, easy-to-handle Jurkat cell-based screening system (Figure 1A).

**Figure 1:**
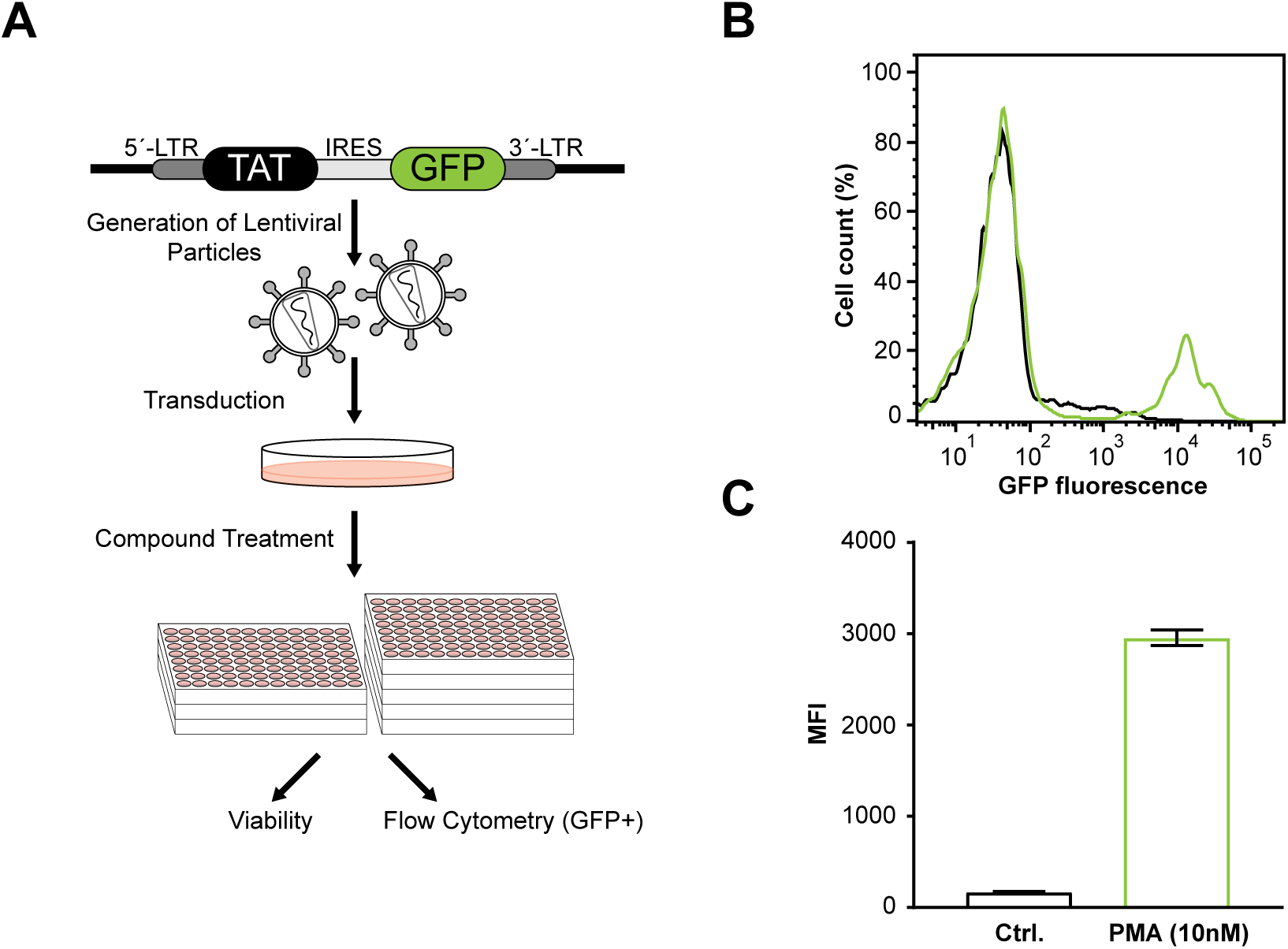
Cell-based reporter system for high throughput screening of LRAs. **A.** Schematic illustration of the screening system. A stable Jurkat-E6 cell-line was created by lentiviral transduction carrying a Tat-IRES-GFP under control of an HIV-1-derived LTR promoter site. Cells were challenged with different compounds in a 96-well based format and subsequently latency reactivation and associated GFP expression was determined by flow cytometry. Simultaneously, cell viability was determined by resazurin assay, to assess compound toxicity. **B.** Histogram of flow cytometry analysis of vehicle-treated control cells (black) and PMA-treated (reactivated) cells (green) **C.** Corresponding bar chart of B, showing the mean fluorescence intensity (MFI) of PMA-treated (positive control) and vehicle-treated (negative control) cells. Error bars represent SD of at least three independent measurements.

Selected candidates were re-tested in a classical latency model at biosafety level 3** conditions for verification. Reactivation of latent HIV by determined lead compounds was confirmed by fluorescence microscopy and lead compounds were applied for *ex vivo* testing of general toxicity in PBMCs as well as of T-cell activation in isolated primary CD4^+^ T-cells to exclude unfavourable side-effects and re-establishment of viral reservoirs upon compound treatment. Our screen identified new potentially effective drugs with significant low toxicity for the treatment of hidden viral reservoirs in HIV-1 infected patients and verified the ability for latency reversal of several already known substances.

## Results

### High troughput screen of epigenetic compounds identifies HIV reversing agents

To allow simplified screening of epigenetic compounds for the ability of HIV latency reversal, we constructed a cell-based reporter system, which enables sample handling under biosafety level (BSL)-1 conditions. For this purpose, we stably transduced Jurkat-E6 cells with a latent reporter HIV‐1 provirus model where a Tat-IRES-GFP reporter cassette is under control of the HIV-1 LTR promoter (27) (Figure 1A).

Positive transduced cells were enriched by fluorescence-activated cell sorting (FACS) and show low basal expression of GFP. However upon stimulation, GFP expression is increased several fold (mean fluorescence intensity; MFI) as shown with the protein kinase C (PKC) activator phorbol 12‐myristate 13‐acetate (PMA), which is known to reactivate provirus via NF-κB and AP-1 signaling (31) and is often used as control compound in reactivation trials (Figures 1B & 1C). Using the MFI values of PMA as positive control and untreated control cells as negative control, we determined the z-factor, which was z = 0.89, verifying the robustness of this assay. Notably, the average z-factor over all measurements in this cell system within this study was 0.77 ± 0.17 (standard deviation). Parallel to reactivation of the GFP reporter, we determined the viability of the cells, to allow detection of toxic compounds/concentrations and to exclude false-positives e.g. due to increased autofluorescence of dead cells. Here, we made use of resazurin (7-Hydroxy-3*H*-phenoxazin-3-one 10-oxide), a non-toxic, cell-permeable and weakly fluorescent phenoxazine dye, which gets reduced by metabolically active and viable cells to the highly fluorescent dye resorufin, thereby providing information about the cellular viability. In addition, qualitative viability data was obtained from cytometry-based forward/sideward scatter information. As all compounds were dissolved in DMSO (10 mM stocks), we first evaluated the background toxicity of DMSO in the reporter cells. DMSO concentrations at or below 1 % resulted in a tolerable toxicity of less than 5 % (data not shown). Thus, highest concentrations of test-compounds in this study were adjusted to 100 µM, being equivalent to 1 % DMSO. Throughout the study, we used 10 nM PMA as positive control and all results were normalized relative to the PMA-based activation. Using the system described, we screened a library consisting of 380 small molecules with biological activity used for epigenetics research. In order to dissect the highest effective and tolerable concentration of each compound, substances were titrated in steps of factor ten from 100 µM to 1 nM. More than 35 % of all compounds (139) lead to a marked reactivation of at least 10% (in comparison to PMA treatment), eight substances caused reactivation of more than 50%. and one compound, the histone methyltransferase inhibitor MI-136 showed an even higher reactivation than PMA (Figures 2A, S2A, S2B). Two other compounds, bisdemethoxycurcumin and the HDAC II inhibitor MC1568, also yielded higher GFP signals than PMA, but were identified in parental Jurkat cells lacking the GFP reporter provirus to have fluorescent properties itself. Notably, with increasing concentrations, an associated toxicity, which additionally often led to certain increase in auto-fluorescence, was observed also for most of the compounds (Figures 2A, S2A, S2B), disqualifying them for further use. This observation was also true for MI-136, where the very high GFP signal was associated with a very high cellular toxicity. To note, treatment with 10 µM PMA itself was associated with a profound reduction of viability up to ~60%, too. A detailed overview of measured percentages of reactivation and cellular toxicity of each compound is given in **Supplemental Figures S2A and S2B**. Overall, we identified in this first round of screening 43 LRA candidates, which showed significant reactivation and at the same time a relatively low toxicity (remaining viability ≥ 75%). Among these small molecules were ~ 35 % HDAC and sirtuin inhibitors, such as vorinostat (zolinza), valproic acid and mocetinostat (MGCD0103) as well as almost as many inhibitors of epigenetic reader domains, like JQ1, I-BET151 (GSK1210151A) and CPI-203.

**Figure 2:**
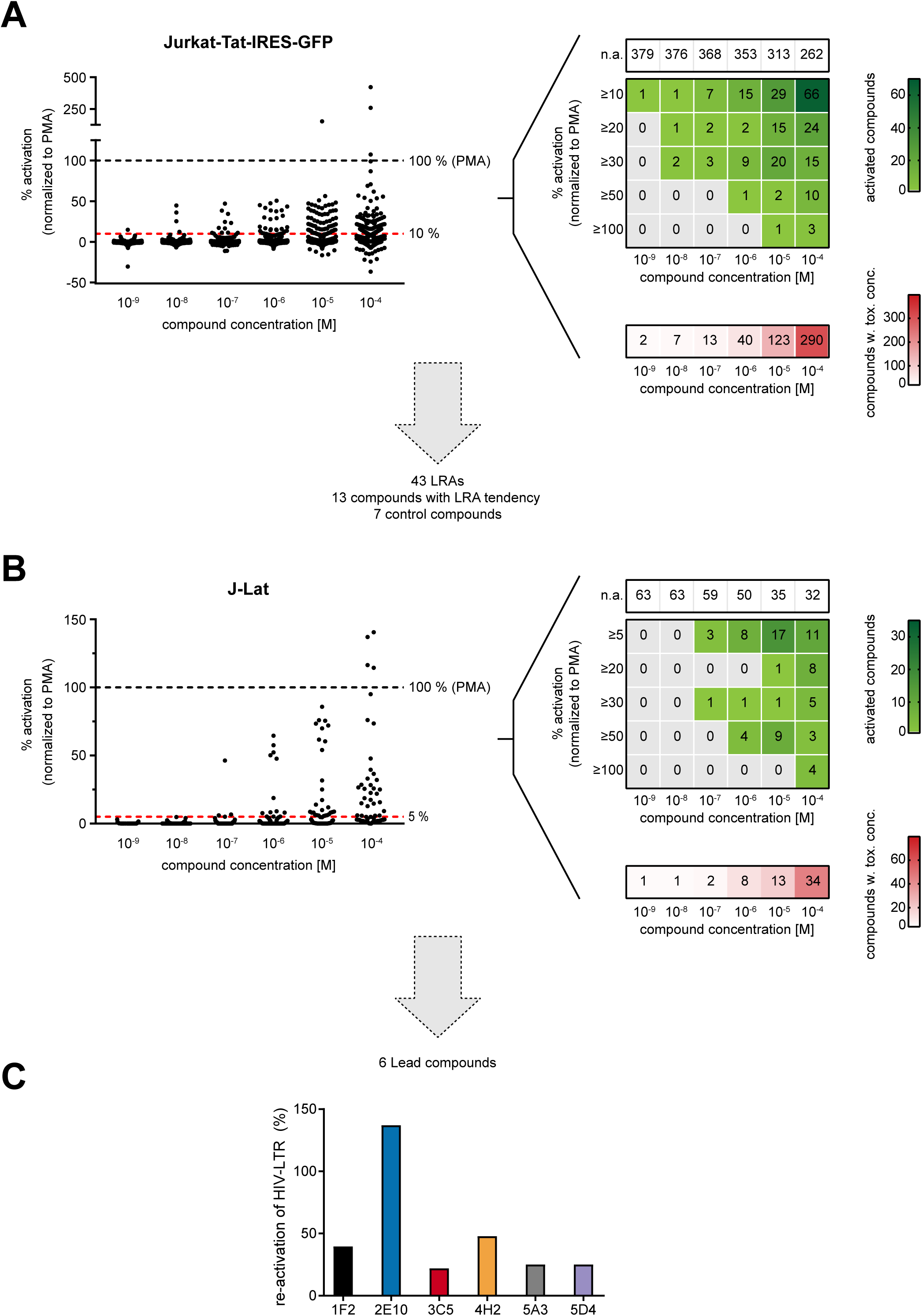
Compound screening. **A.** Primary screen based on Jurkat-E6 reporter cells. Left graph displays the average percentage of activated cells for each compound and concentration normalized to PMA-treated control cells. Activation threshold was set to 10 % and only compounds causing ≥ 10 % reactivation were defined as potential LRA. Corresponding upper heatmap on the right displays the number of compounds at the indicated concentration leading to a reactivation of ≥ 10 %, ≥ 20 %, ≥ 30 %, ≥ 50 %, or ≥ 100 % in comparison to PMA treatment. n.a. (no activation, threshold < 10 %). The small heatmap (lower right site) displays the amount of compounds at the indicated concentration leading to non-tolerable cellular toxicity (viability threshold: 75 %). **B.** Secondary screen of dedicated compounds identified in A. Screen was performed in J-Lat cells. Graphs essentially as described in A, but with threshold for activation of 5 %. **C.** Bar chart displays mean percentage of reactivation in J-Lat cells caused by challenging the cells with the indicated six lead compounds.

### Validation screen of selected compounds in latency model cells

The 43 candidates from the high throughput screening in Jurkat-E6 cells were further validated under biosafety level BSL3** conditions using J-LAT cells, a well-known latency model system, also derived from Jurkat T-cells that had been transduced with near full-length, Δ*env* and *nef*-defective HIV R7, that carries a *gfp*-reporter cassette in the *nef* locus (27). Another 13 compounds, including the HDAC inhibitor SR-4370 or the methyltransferase inhibitor AMI-1, were included as they showed either reactivation very close to the 10 % threshold or a distinct reactivation but a toxicity close to the threshold of 75 % viable cells. These compounds are classified hereinafter as “borderline”. Furthermore, we added 7 compounds for control reasons that showed no or very little reactivation in the Jurkat-LTR-GFP system (Figures 2B, S3). Data in the J-Lat system mirrored well the results from our first screen. While none of the used control substances was able to reactivate the viral promoter, most of the LRAs as well as the “borderline”-compounds from the first screen showed a substantial reactivation ability. Moreover, the signal background in the J-Lat model system was lower, allowing us to set the threshold for reactivation to 5% GFP fluorescence (compared again to 10 nM PMA as calibrator substance). However, cellular toxicity and signal intensity varied in both cell systems. For example, the average viability at the highest concentration of 100 µM significantly shifted from 53.7 % as determined in our pilot screen to 70.5 % in the J-Lat model system (p= 0.0002; one-way Anova). At the same time, the necessary concentration for LTR-activation was increased for most of the compounds when compared to the Jurkat-LTR-GFP model. Based on their reactivation efficiency and toxicity, we identified six lead compounds from the second screen, which showed considerable induction of the HIV-LTR promoter (Figures 2C). The observed latency reactivation was also further validated by fluorescence microscopy, confirming the flow cytometry results (Figure 3A).

**Figure 3:**
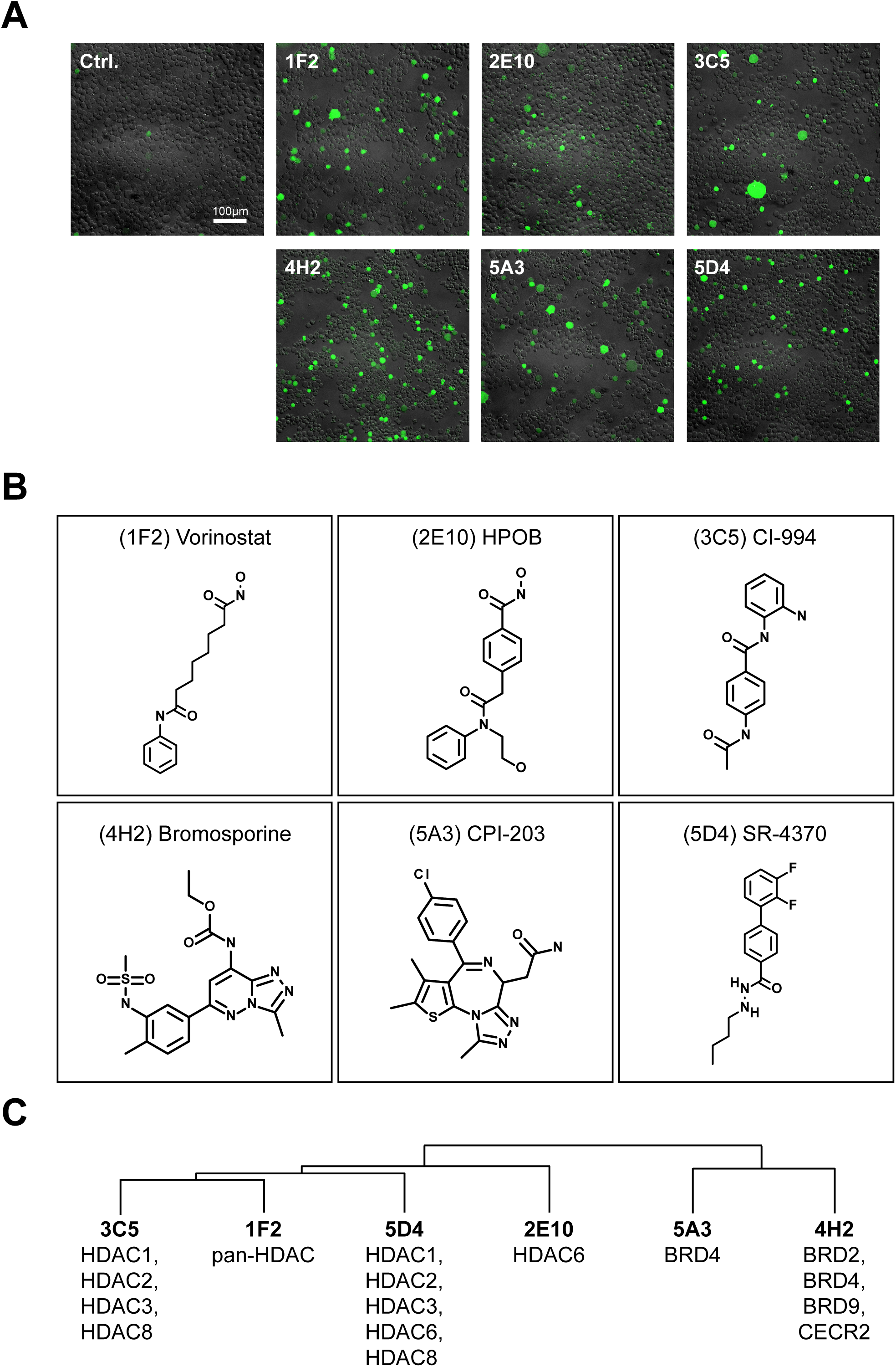
Verification of lead compound function and physicochemical analysis. **A.** Complementary analysis of latency reversing function of the identified lead compounds in J-Lat cells by fluorescence microscopy. Representative pictures of J-Lat cells challenged for 24 h either with the indicated compound or with vehicle only (control). **B.** Chemical structure of the lead compounds and common name. **C.** Hierarchical clustering of lead compounds based on physicochemical properties derived from the compound structure. Known targets of each compound are depicted below the compound number. Clustering was performed with ChemMine Tools (https://chemminetools.ucr.edu/).

The six LRAs comprised the HDAC inhibitors vorinostat (SAHA), CI-994 (Tacedinaline), SR-4370, and HPOB as well as the two Bromodomain (BRD) inhibitors CPI-203 and bromosporine. While, several HDAC inhibitors, including the identified SAHA and both BRD inhibitors have been already decribed for reactivation of HIV latency (32–34), the class I HDAC inhibitor SR-4370 as well as the HDAC6 inhibitor HPOB have not been tested so far. Here, especially HPOB draw our attention as treatment resulted in a maximum response of 137 % compared to PMA (Figure 2C), while the related toxicity was still low (82 % viability). Analysis of physicochemical features of the compounds via ChemmineR (30) and subsequent clustering revealed diverse chemical structures (Figure 3B, 3C) and was in good agreement with known targets of the compounds separating BRD and HDAC inhibitors. Notably, within the group of HDAC inhibitors both newly identified drugs clustered relatively close together, being also reflected by their affinity for HDAC6.

### Determination of T-cell activation and side-effects on other leucocytes

Compound induced reactivation of latent HIV may not only initiate viral gene transcription but might also overlap with global T-cell activation, which is associated with unfavorable cytokine production (35) and could enhance *de novo* infection of additional cells. Therefore, an important ability of LRAs for clinical use is a significant reactivation capability without T-cell activation. To test possible T-cell activation properties of the lead compounds, we isolated resting CD4^+^ T-cells from PBMCs of anonymized healthy donors and determined the surface expression of the early (CD69) and late activation antigens (CD25) by fluorescence-based flow cytometry 24 h post treatment. Here, vehicle-treated cells served as negative control and PHA or CD3/CD28-bead stimulated cells were used for positive control (Figure 4). None of the six compounds showed an increase in CD25, CD69, and CD25/CD69 positive cells, comparable to vehicle-treated control cells, while activation by PHA or CD3/CD28 magnetic beads caused significant upregulation of both activation markers.

**Figure 4:**
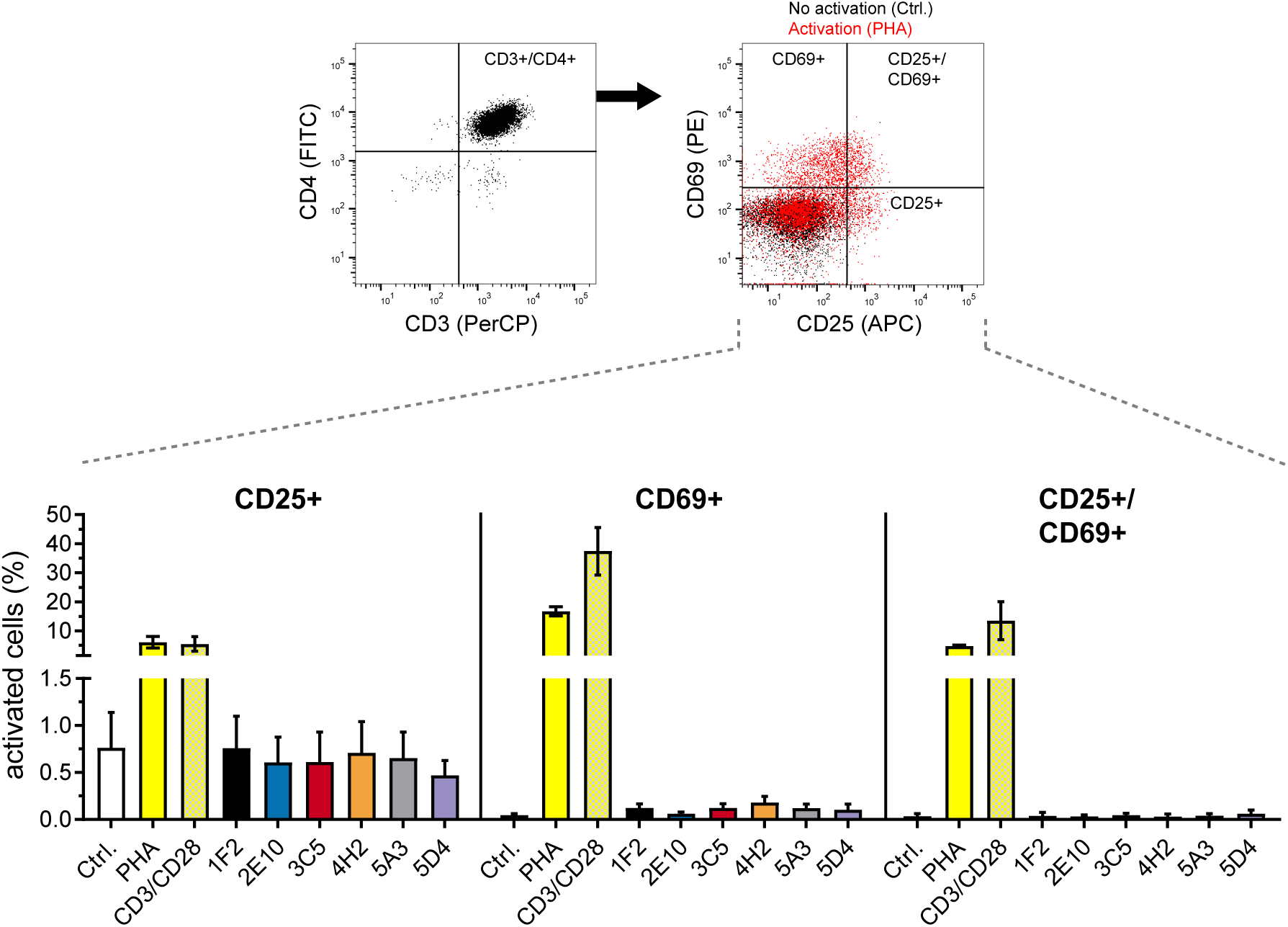
CD4^+^ T-cell activation. Resting CD4^+^ T-cells were isolated from blood of healthy donors and challenged for 24 h with either PHA or CD3/CD28 beads (positive controls) or with the respective compound. Vehicle-treated cells were used as negative control (control). Subsequently, cells were stained for CD3, CD4, CD25 and CD69 and analyzed by flow cytometry for increased expression of activation markers CD25 and CD69. Flow cytometry plots show gating for CD3+/CD4+ subset (left) and corresponding flow cytometry plot (right) shows distribution of CD25/CD69 on control cells (black) and activated, PHA-treated cells (red). Bar charts indicate amount of CD25, CD69, and CD25/CD69 positive cells. Error bars represent SD form three different donors.

Another obstacle for clinical use of LRAs might be cytotoxic side-effects of these compounds on other immune cells, which would affect the efficacy of therapy. Thus, in addition to the T-cell activation test, we performed flow cytometric immunophenotyping of PBMCs 24 h post compound treatment to exclude unfavorable toxicity to leukocytes in general (Figure 5). We observed no difference in global cell viability (as measured by 7-AAD staining) nor significant changes in the amount of cells expressing CD45^+^, CD3^+^, CD4^+^, CD8^+^, CD14^+^, CD19^+^, CD16/56^+^, CD15/56^+^/CD3^+^ surface antigens. However, a higher but not significant fluctuation was observed in case of CD14^+^ monocytes, which might be associated with induced proliferation/differentiation of myeloid cells (36).

**Figure 5:**
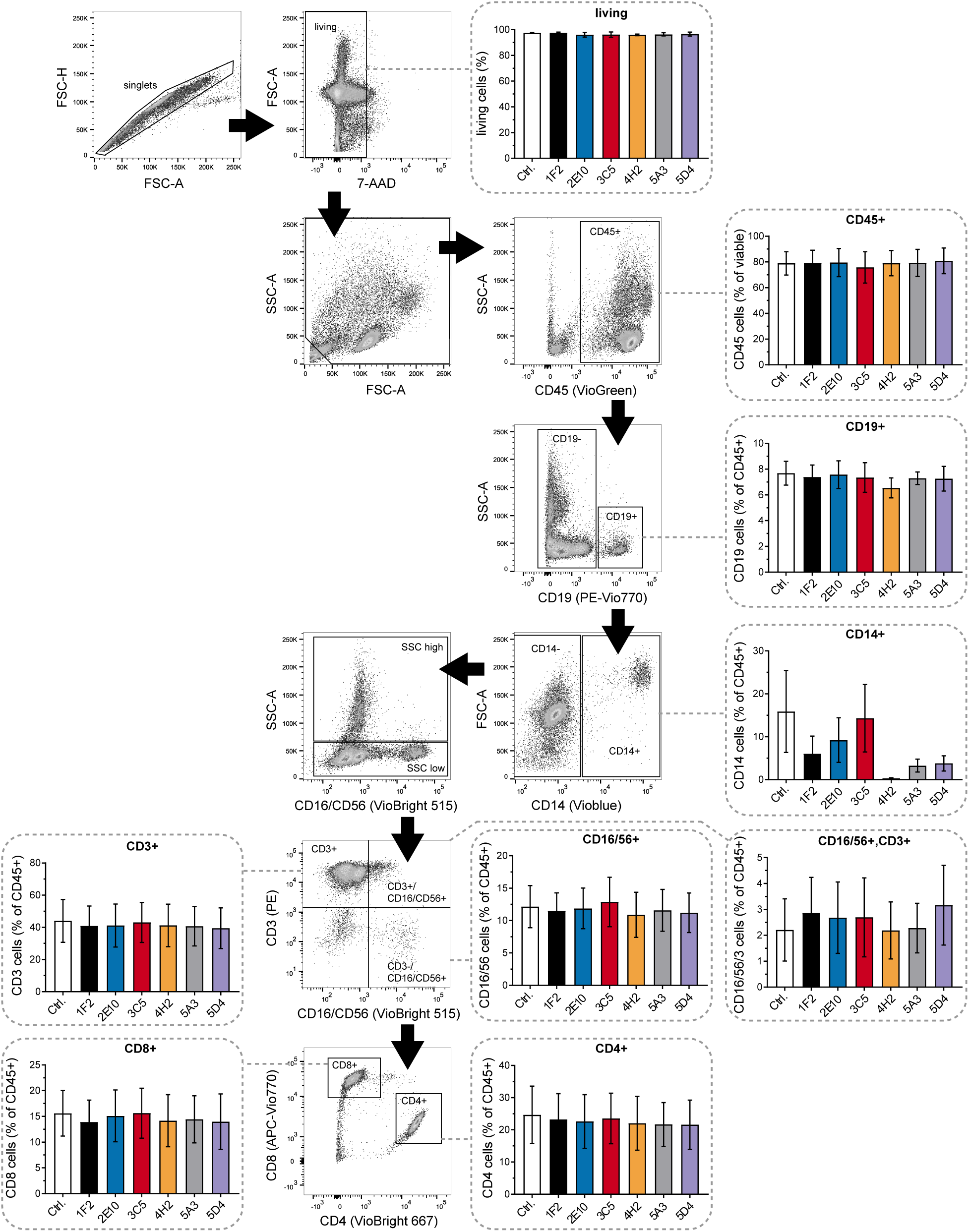
Immunophenotyping of PBMCs. PBMCs, derived from healthy donors, were challenged for 24 h with the respective compound or with vehicle control and subsequently analyzed by flow cytometry for certain cellular markers. The flow cytometry plots show representative data of control cells and the underlying gating strategy. Cells were first gated for singlets, tested for viability via 7-AAD stain, and cell debris was removed by classical SSC/FSC analysis. Derived living single cells were analyzed for CD45 expression as general leucocyte marker and leucocytes were further discriminated in B-cells (CD19) and monocytes (CD14). CD19/CD14 negative cells were than subdivided into eosinophils/granulocytes (SSC high) and CD16dim SSC low fraction. CD3 and CD56 were used for determination of CD3+ T-cells, CD56+ NK cells and CD3+/CD56+ T-cell like NK-cells. Subsequently, CD3+ cells were analyzed for CD4+ and CD8+ T-cells. Associated data for each cell type is presented as bar chart diagram and shows results from three different donors. Error bars represent SD.

In conclusion, our data demonstrate the importance of acetylation-related events during latency, confirm earlier findings, and identified two new compounds, HPOB and SR-4370, which are promising candidates to optimize common treatment strategies. Both compounds might help to target persistent latent viral reservoirs in HIV-1 infected patients.

## Discussion

Current ART is highly effective in the reduction of viral loads to undetectable levels in plasma and allows a largely normal life of infected individuals. However, it does not eradicate latent cellular reservoirs of HIV, making lifelong therapy necessary. Moreover, studies have shown that, despite effective ART, viral replication can occur in certain sanctuaries of the lymphoid tissue, where the concentration of therapeutics is not sufficient to block viral replication (10, 12, 37). Thus, several trials have been undertaken to deliver on the one hand drugs more effectively to the viral replication sites and on the other hand to identify and destroy established latent reservoirs, with the aim of full viral clearance. Recently, especially the cellular mechanisms maintaining the latent state are attempted to be overcome by pharmaceutical interventions, which consequently are thought to reactivate those cells and make them accessible for immune cells and common antiretroviral drugs.

However, so far trials have not been clinically successful yet. Thus, the discovery of either cellular factors that maintain latency or novel latency-reversal agents is highly demanded and anticipated to support current “shock‐and‐kill” approaches, which are still considered to be the clinically most feasible method.

Consistent with this aim, and based on recent findings showing the importance of posttranslational modifications (such as lysine-acetylation) for viral replication and latency (11), the idea of this study was to screen for novel latency-reversing agents, focusing in particular on epigenetic-active drugs. For this, we constructed an easy-to-use reporter cell line, which allows handling under BSL-1 conditions (cells were verified to be free of residual viral particles) and thus qualifies for less costly HTS assays. Initial screening in this cell model, comprising compound titrations and toxicity studies, identified around 10 % of all tested small molecules within this study to be able to stimulate the viral LTR-promoter in a dose dependent manner. To ensure validity of results and to compare common cell models, we verified the compound-induced reactivation in J-Lat cells, an often used latency model cell line, under BSL3** criteria. This second screen mainly confirmed our data from the first screen and revealed six lead compounds that showed on the one hand very good reactivation properties and on the other hand low toxic effects in both cell systems. Interestingly, we observed that, while most of the lead compounds reactivated the LTR in individual cells accompanied with high GFP expression as shown by microscopy (Figure 3A), HPOB (compound 2E10) showed a more equally distributed GFP expression, although with lower fluorescence intensity. Thus, HPOB may reactivate a wider range of viral reservoirs, than common class I HDAC inhibitors.

Physicochemical analysis of the lead compounds and literature research, clearly underlines the importance of lysine-acetylation in general for maintenance of HIV-related latency as all six compounds could be assigned either to histone deacetylase (HDAC) inhibitors or inhibitors of bromodomain (BRD)-carrying proteins, which serve as “readers” of lysine-acetylations. Molecules like vorinostat and CI-994, which have been described already in earlier studies as potent LRAs and have been investigated in clinical trials (32, 38), targeting mainly class I HDACs, which have been shown to be associated with histone-acetylation and e.g. the transcriptional repressors YY1 and LSF (39–42). Likewise, the two identified inhibitors of BRD containing proteins, bromosporine and CPI-203, which target amongst others BRD4, which has been associated with the repressive SWI/SNF chromatin remodeling machinery (43), have been shown to efficiently reverse latency (33, 34). Interestingly, both within this study newly discovered LRAs, SR-4370 and HPOB, target contrarily to the other four compounds, also the cytosolic deacetylase HDAC6, which has been found to be associated with antiviral activity against HIV by induction of autophagic degradation of the HIV Vif protein (44) as well as by tubulin-associated prevention of HIV-1 envelope-dependent cell fusion and infection (45). In addition, HDAC6 has been shown to interact with and to deacetylate the viral trans-activator of transcription (Tat), resulting in the suppression of Tat‐mediated transactivation of the HIV‐1 promoter (46). Because of these HDAC6-related functions, and our screening results (which include also other effective reversing HDAC6 inhibitors like tubastatin A HCl), further studies should be undertaken towards the role of HDAC6 in viral latency.

Another important observation from this study is the general compatibility of the identified LRAs with other immune cells such as cytotoxic T-cells or cells of the myeloid lineage, which was examined using primary cells. First, compounds did not activate resting CD4^+^ T cells as determined by the activation-markers CD25 and CD69. This is a prerequisite for further clinical trials as T-cell activation might be accompanied with cytotoxic cytokine release (47) and involves the risk of increased susceptibility of neighboring T-cells and thus the associated creation or relocation of latent reservoirs. Second, none of the six compounds showed an unfavorable impact on any leucocyte-associated cell type as the number of B-cells (CD19^+^), monocytes (CD14^+^), NK (CD16/56^+^) and NK-like T-cells (CD16/56^+^ and CD3^+^) as well as of CD4^+^ and CD8^+^ T–cells did not change upon treatment in comparison to untreated control cells. Third, and very important, no increased overall toxicity of the PBMC population was discovered, underlining the potential usability of the compounds. Furthermore, all lead compounds were able to induce profound reactivation of viral transcription in different cell systems. As the different systems most likely also resemble different areas of integration as well as variances in the epigenetic landscape, it can be assumed that the compounds identified in this study will thus also show a broad applicability within the clinical setting.

In conclusion, the described simplified screening for latency reversing agents in combination with an epigenetic-active drug library verified former studies and revealed two novel compounds, which may be used to induce HIV transcription in a broad range of latent viral reservoirs. We suggest to include both compounds in future clinical studies to assess their clinical usability.

## Material and methods

### General reagents and chemical compounds

All standard reagents and chemical compounds were obtained either from Carl Roth GmbH + Co. KG/Germany, Sigma Aldrich/Germany, Thermo Fisher Scientific/USA, or TH. Geyer/Germany if not otherwise stated. The epigenetics compound library (catalog # L1200) has been derived from TargetMol/USA.

### Cell culture

Jurkat-E6, J-Lat (clone 8.4), and Jurkat-E6-LTR-Tat-EGFP cell lines were grown in RPMI 1640 medium (Gibco) supplemented with 10 % FCS and 100 U/ml penicillin, 100 µg/ml streptomycin at 37°C under a 95 % air / 5 % CO_2_ atmosphere. Hek293T cells were grown under the same conditions on DMEM medium (Gibco). All cells were confirmed to be mycoplasma negative (Lonza MycoAlert).

### Production of lentiviral particles and transduction of Jurkat-E6 cells

For generation of lentiviral particles, 15 × 10^6^ Hek293T cells were co-transfected with 7 µg pCMV.d8.9 (packaging vector), 8 µg pMD2.G (envelope vector coding for vesicular stomatitis virus glycoprotein VSV-G), and 11 µg pEV731 plasmid (transfer vector containing the LTR-Tat-IRES-GFP sequence (27)) using polyethylenimine procedure (28) with a DNA to PEI ratio of 1:3. 48 h post transfection, virus containing supernatant was harvested and filtered (0.45 μm, Millex-HV Filter; Millipore). Subsequently, the filtrate was purified and concentrated for viral particles by sucrose-cushion-based ultracentrifugation. For this, 25 % sucrose (dissolved in H_2_O) was overlaid with filtered supernatant and centrifuged in a swing-out rotor ((SW 32 Ti; Beckman Coulter or SureSpin™ 630/36; Thermo Fisher Scientific) at 110,000 × g and 4°C for 90 min. After centrifugation, pelleted viral particles were resuspended in PBS and stored at −80°C. Concentrated virus stocks were titrated on Jurkat-E6 cells and cells were cultured at 37°C and 5% CO2. Cells were verified to be free of residual viral particles 10 days post transduction using the QuickTiter™ Lentivirus-Associated HIV p24 titer kit (Cell BioLabs Inc.).

### Cell sorting of reporter cell line

Transduced Jurkat-E6 cells were stained with SYTOX™ Blue dead cell stain (Invitrogen) according to manufacturer’s instruction and sorted for viable, GFP-positive and -negative cells using a FACSAriaFusion flow cytometry sorter (BD). Sorting was performed at the Core Facility Flow Cytometry at Biomedical Center Munich of the LMU Munich. Post sorting, GFP-positive cells were cultured for 48 h in RPMI medium supplemented with 30% FCS and 100 U/ml penicillin, 100 µg/ml streptomycin until replacement with standard medium to reduce sorting-related mortality.

### Compound screening

Compounds from the epigenetics compound library were titrated (1 nM to 100 µM final concentration) on 2×10^5^ Jurkat-E6-LTR-Tat-EGFP cells per well and cells were cultured for 24 h. Subsequently, cells were washed with PBS and EGFP fluorescence was analysed in a FACSLyric flow cytometer (BD Biosciences). Untreated (Vehicle-treated) and PMA-treated (10 nM) cells were used for control. Quality of screen was monitored for all titration experiments by analysis of the Z-factor (29) according to the following formula:

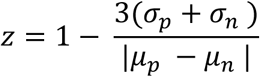

with standard deviation (σ) of positive (p) and negative (n) control and the means (µ) of the positive (p) and negative (n) control. For validation of selected compounds, screening was repeated in J-Lat cells at biosafety level S3**. In this case, cells were not only washed with PBS, but also inactivated for 90 min with 4 % PFA dissolved in PBS prior to flow cytometric analysis.

### Viability assay

Viability of Jurkat-E6-LTR-Tat-EGFP cells and cytotoxicity of compounds, respectively, was analyzed 24 h post treatment using alamarBlue™ Cell Viability Reagent (Invitrogen) according to manufacturer’s instruction. AlamarBlue was added to cells 4 h prior analysis and fluorescence of formed resafurin was measured in black 96-microwell flat-bottom plates (Brand) in a CLARIOstar Plus microplate reader (BMG Labtech). Untreated, and PMA-treated (10 nM) cells served as controls. Additionally, viability was controlled by flow cytometry using forward/sideward scatter method.

### Microscopy of compound-treated J-Lat cells

J-Lat cells were freed 24 h post compound treatment from cell debris by low speed centrifugation (23 × g, 5 min, RT) and afterwards harvested by centrifugation (400 × g, 5 min, RT). Cell pellets were re-suspended in culture medium supplemented with 4 % PFA (Electron Microscopy Sciences) and inactivated for 90 min at RT. Subsequently, cells were pelleted (1200 × *g*, 10 min, 4 °C), washed with PBS, and embedded in Prolong™ Diamond Antifade Mountant (Invitrogen) on microscopy glass slides sealed by a glass coverslip. Samples were imaged (brightfield exposure time: 200 ms, GFP exposure time: 100 m) using the Eclipse Ti2 microscope (Nikon) in combination with a DS-Qi2 camera (Nikon) under control of NIS Element AR software (v. 5.0, Nikon). Acquired images with 0.365 µm/pixel resolution were overlayed and analyzed using ImageJ (v. 1.52a) and the ND2 Reader plug-in (Nikon Instruments Inc).

### CD4^+^ T-cell activation assay

Blood cones for isolation of CD4^+^ T-cells were derived from the Terumo BCT leukocyte reduction system used at the Hospital of the University of Munich, Dept. of Immunohematology, infection screening and blood bank (ATMZH). Content was diluted with PBS (Gibco) and CD4^+^ T-cells were isolated via the Easy-Sep™ Rosette Human CD4^+^ T-cell enrichment kit (Stemcell Technologies, Canada) according to manufacturer’s instructions and purity of isolated CD4^+^ T-cells was controlled on a FACSverse™ flow cytometer (BD Biosciences) using BD Tritest CD4/CD8/CD3 antibody stain (BD Biosciences). Derived non-activated (resting) T-cells were adjusted to 2×10^6^ cells/ml in RPMI medium, supplemented with 10% FCS and 100 U/ml penicillin, 100 µg/ml streptomycin. For activation testing, 2×10^6^ cells were cultivated for 24 h in presence of either Human T-Activator CD3/CD28 dynabeads (Thermo Fisher Scientific), according to manufacture’s instruction, or 10µM phorbol-12-myristate 13-acetate (PMA; Sigma Aldrich), or the respective compounds from the compound library. Subsequently, activation of T-cells was measured by flow cytometry using the anti-human CD4 (FITC)/CD69 (PE)/CD3 (PerCP) antibody staining solution (BD Biosciences) in combination with anti CD25 (APC) (BD Biosciences). Vehicle-treated (resting) T-cells served as control.

### Immunophenotyping

PBMCs were derived from blood cones (see T-cell activation assay) by Ficoll (Biochrom) density gradient centrifugation. PBMCs were washed three times with PBS (Gibco) and adjusted to 2×10^6^ cells/ml RPMI medium supplemented with 10% FCS and 100 U/ml penicillin, 100 µg/ml streptomycin. Cells were treated with the respective compounds from the compound library or vehicle-control and cultivated for 24 h. Subsequently, cells were washed twice with cold PBS and stained with the human 8-color immunophenotyping kit (Miltenyi), according to manufacturer’s procedures. Stained cells were analyzed in a FACSLyric flow cytometer (BD Biosciences). First, doublets were eliminated by gating for single cells in forward scatter area (FSC-A) versus forward scatter height (FSC-H). Viable cells were identified by 7-AAD staining and after removal of residual cell debris, the amount of CD45+ cells was determined. CD45+ leucocytes were further separated into CD14+ and CD19+ cells and remaining cells were divided by site scatter into SSC high (neutrophils, eosinophils) and SSC low cells. The latter were further divided into CD3+, CD16/56+, and CD16/56+/CD3+ cells. Finally, CD3+ cells were separated into CD4+ and CD8+ cells.

### Data analysis

Flow cytometry data analysis was performed using FlowJo software (TreeStar, v. 10) and further data analysis was performed using Microsoft Excel (v. 2016) and GraphPad prism software (v. 7.05). Physicochemical analysis of lead compounds was performed using ChemmineR (30).

### Ethics statement

All work with J-Lat cells was performed in a BSL3** facility. Work with HIV has been approved by the Bavarian government (AZ 55.1GT-8791.GT-2-1206-11). Usage of blood cones was approved by the ethics committee of the LMU München, Munich, Germany with the project No.: 17-202-UE.

## Acknowledgements

pMD2.G was derived via addgene (www. addgene.org) from Didier Trono and pEV731 was kindly provided Eric Verdin. J-Lat cells were obtained through the NIH AIDS Reagent Program from Eric Verdin (27). We would like to thank Rebecca Engels (AG Sewald; MvPI Munich/Germany) for technical assistance.

## Author contributions

FS established the Jurkat-LTR-GFP model cell line. AZ, LC, FS, and CS designed and performed high-throughput screenings. LC performed microscopy experiments. CS and AZ performed data analysis. All authors discussed the results, provided critical feedback and contributed to the final manuscript.

